# Exploring phenotypic and genetic variation in *Lactuca* with GWAS in *L. sativa* and *L. serriola*

**DOI:** 10.1101/2025.06.27.661939

**Authors:** Sarah Mehrem, Guido van den Ackerveken, Basten L. Snoek

## Abstract

Crop wild relatives provide valuable insights into trait diversity and the genetic basis of agronomic traits. In the genus *Lactuca*, domesticated lettuce (*Lactuca sativa*) and its wild progenitor, *Lactuca serriola*, have been extensively studied, yet broader wild species remain underrepresented. Here, we present a phenotypic dataset of 550 *Lactuca* accessions, including 20 wild relatives, capturing plant morphology, pigmentation, and pathogen resistance traits derived from images and genetic resource collections. To investigate the genetic basis of these traits, we used a jointly processed SNP set for *L. sativa* and *L. serriola*, applying an iterative two-step GWAS approach, enabling the dissection of multiple loci per trait. We identified both known and novel QTLs associated with anthocyanin accumulation, leaf morphology, and pathogen resistance in *L. sativa* and *L. serriola*. Importantly, we identified *L. serriola*-specific QTLs undetected in *L. sativa*, revealing unique genetic architectures underlying anthocyanin biosynthesis and leaf morphology in the wild progenitor. These findings expand the knowledge of *Lactuca* beyond cultivated varieties, highlighting the potential of wild species for breeding applications. Our dataset and results provide a foundation for further investigations into the evolutionary and agronomic significance of *Lactuca* diversity.

## Introduction

Plant phenotyping and genomics are essential for understanding the genetic basis of plant traits, thereby advancing crop improvement. The inclusion of crop wild relatives (CWR) in genetic studies has enhanced our understanding of agronomic trait development, domestication, and evolutionary history. Investigating CWR provides valuable insights into the genetic mechanisms that underly domestication traits, as demonstrated in studies on various crops [1–6]. Studies on wild relatives of tomato (*Solanum lycopersicum*) and potato (*Solanum tuberosum*) have revealed that domestication traits, such as fruit morphology and fertility, originated in semi-domesticated or wild populations, and were later introduced into cultivated varieties [1, 3]. These findings highlight the importance of CWR as reservoirs of genetic diversity for enhancing existing or introducing novel traits.

The *Lactuca* genus of the *Asteraceae* family encompasses over 100 species, including both wild and domesticated taxa, with lettuce (*Lactuca sativa*) being the most widely cultivated species [7]. After its initial domestication around 4000 BC, lettuce was first cultivated as an oilseed crop. Over time, it has undergone significant diversification, resulting in a wide range of cultivars with diverse leaf shapes, colors, textures, and growth forms. These include loose-leaf varieties (Cutting-type), heading types (Crisp, Butterhead, Cos), as well as stem-type cultivars. The diversity within lettuce makes it a valuable model for studying the genetic basis of agronomic traits in leafy greens, including morphology, stress resistance, and development [8].

Prickly lettuce (*Lactuca serriola*) is considered the wild progenitor of lettuce [9]. Along with *L. virosa* and *L. saligna*, it continues to be one of the primary wild *Lactuca* species used for breeding new lettuce varieties [10]. To date 20 *Lactuca* species are considered members of the lettuce gene pool [8, 11–13]. Included in the primary gene pool are the main species *L. sativa* and *L. serriola*, with further members being *L. aculeata* and *L. altaica*. The secondary gene pool consists of *L. saligna* alone. The tertiary gene pool includes *L. virosa*, *L. georgica, L. quercina*, *L. sibirica*, *L. tatarica*, and *L. viminea*. Several other related species outside of these gene pools, such as *L. indica*, *L. homblei*, *L. raddeana*, *L. canadensis*, *L. biennis*, *L. floridana*, *L. tenerrima*, *L. palmensins* and *L. perennis* contribute to the diversity of traits explored in this study. Despite the importance of wild *Lactuca* species for breeding lettuce, they have generally been underrepresented in genetic resource collections, except for *L. serriola*, *L. virosa* and *L. saligna* [8, 14].

Genome-wide association studies (GWAS) and recombinant inbred line (RIL) mapping in *L. sativa*, and to a lesser extent in *L. serriola*, have provided valuable insights into the genetics underlying trait variation, including disease resistance to *Bremia lactucae*, plant architecture and pigmentation, shelf life, heading, seed coat color, and metabolite content [8, 15–20]. Moreover, most of these studies have focused on the phenotypic variation in *L. sativa* and *L. serriola*. While *L. sativa* has received extensive attention due to its commercial relevance, *L. serriola* remains less explored. This lack of attention applies even more to the other wild relatives, *L. virosa* and *L. saligna*, where, aside from a few studies mapping pathogen resistance traits, virtually no QTL mapping has been performed [8, 21–25].

In this study we curated a broad phenotype collection that includes *L. sativa*, *L. serriola*, *L. virosa*, *L. saligna* and 17 additional CWR. The phenotypes were curated from both existing germplasm collections, as well as images of adult and juvenile plants, flowers, and seeds. This collection includes phenotypes from different growth stages of the same accessions, and highlights under-represented wild *Lactuca* species, which are rarely documented together. For future sequencing studies with a more extensive range of wild *Lactuca* species, this collection provides a strong basis for investigating trait diversity beyond *L. sativa* and *L. serriola*. With SNP-based genotype data for both *L. sativa* and *L. serriola* accessions, we extend the results of multiple GWAS and QTL mappings previously conducted in both species, providing one of the largest collections of QTLs for these species. Furthermore, we perform GWAS as a two-step process, where the most significant SNP of the first GWAS iteration is added as a fixed effect to the second GWAS iteration. This method allowed us to resolve multiple QTLs per trait, including previously undetected QTLs. By uncovering additional associations, we highlight possible new breeding targets for traits related to plant morphology, pigmentation, and pathogen resistance. Overall, our approach offers deeper insights into trait diversity, particularly within wild species.

## Methods

### Curating phenotypes of *Lactuca* accessions

We collected 135 phenotypes for 550 *Lactuca* accessions. The phenotypes are represented using three data types: binary, ordinal, and quantitative. We obtained photos of seeds, flowers, juvenile and adult plants through the Center for Genetic Resources Netherlands (CGN), including pictures from a special collection (http://www.wur.eu/cgnsc002) [26]. From these photos the following phenotypes were scored according to the lettuce calibration book: Leaf blade: division, leaf: anthocyanin coloration, leaf: intensity of anthocyanin coloration, leaf: type of anthocyanin distribution, leaf: blistering, leaf blade: depth of incisions on margin on apical part, leaf blade: degree of undulation of margin [27]. Furthermore, trait leaf blade: type of incisions on apical part was scored according to the manual, however on all plants, not only on plants that had shallow incision margins, as stated in the manual. The scoring of flower related phenotypes was done by eye. The acquisition of remaining phenotypes scored from these pictures is described in supplemental table S1. In addition, we acquired further phenotypic data for all 550 accessions through the CGN lettuce collection [28].

### Analysis of phenotypic variation

Phenotypes were summarized into six broader categories: Pigmentation, which includes all leaf-related color traits; Seed, encompassing all seed-related traits; Pathogen Resistance, covering traits associated with pathogen responses; Morphology, describing traits related to leaf shape and plant structure; Flower, which includes all flower-related traits. All further analysis was performed using R (Version 4.2.2) [29]. The cor() function of base R with the parameters “use=pairwise” was used to calculate Spearman’s coefficient for quantitative and ordinal phenotypes. For correlation we only considered phenotypes with less than 70% of values missing. We describe phenotypic (dis)similarity between all *Lactuca* accessions using Gower’s distance with the daisy function with metric=”gower” of the R-package cluster (Version 2.1.4) [30]. For computing Gower’s distance, we included phenotypes with less than 50% of data points missing and adjusted for over-representation of morphological traits by adding weights based on frequency of phenotype group. Heatmaps for correlation and distance matrices were created using the R-package ComplexHeatmaps (Version 2.14.0) [31].

### *Lactuca* sp. tree

The tree depicting *Lactuca* sp. taxonomy was curated manually from multiple phylogenies to place all *Lactuca* species that are included in this study. For the basis, we used a coalescence-based phylogeny of single-loci nuclear genes of 12 *Lactuca* species [8]. *L. dregeana* was removed from the phylogeny because there was no phenotypic data available. The following species were added based on nuclear ribosomal DNA phylogenies [32]: *L. quercina*, *L. raddeana*, *L. biennis*, *L. floridana*, *L. tenerrima*, *L. sibirica*, *L. tartarica*, *L. viminea* and *L. perennis*. *L. canadensis*, was placed into the tree, based on nuclear ribosomal DNA internal transcribed spacer phylogenies [33]. The *Lactuca* tree was then annotated based on reported germplasm pools that underly genetic variation of *L. sativa* [8]. *L. georgica* was changed to the secondary germplasm pool based on [34]. The tree was further annotated into two clades [32]. For visualizing the tree, the R package ggtree (Version 3.6.2) was used [35]. The tree is included in the supplement in Newick format (**Lactuca_tree**).

### SNP matrix

The SNP data for *L. sativa* and *L. serriola* accessions included in this study was previously published by Dijkhuizen *et al*. (2024), where the methodology for sequencing, SNP calling and SNP filtering is described [36, 37]. In short, SNPs were called against the Salinas V8 Lettuce reference genome, downloaded from NCBI (GCF_002870075.4, https://www.ncbi.nlm.nih.gov/genome/352) using the GATK HaplotypeCaller embedded in the Nextflow Illumina Analysis Pipeline (https://github.com/UMCUGenetics/NF-IAP/) [38]. Only bi-allelic SNPs were kept for further analysis. One SNP matrix per species was generated, using the same filtering and clumping parameters. SNPs were clumped in batches of 2000 neighboring SNPs, with a clumping criterion of differing in less than 5 accessions. The SNP with the highest MAF was chosen as the SNP clump representative. SNP alleles were coded with 0 for homozygous reference allele, 1 for heterozygous and 2 for homozygous alternative allele.

### GWAS

GWAS was performed per species. Per phenotype SNPs with a MAF < 5% were excluded. Binary phenotypes were represented as binary arrays, ordinal and quantitative phenotypes as numerical. Binary and ordinal phenotypes were excluded if more than 95% of accessions had the same score. Phenotypes with less than 80 observations were excluded. GWAS was performed in R (Version 4.2.2) using the lme4QTL and pbmcapply packages with the script Run_GWAS.R [39, 40]. *L. sativa* accessions LK086, LK196, LK197, LK198, LK199, LK200, and *L. serriola* accessions LK203, LK204, LK206, LK208 were excluded from GWAS due to being recorded as an oilseed variety or *L. sativa* x *L. serriola* cross, reported as sample switches before sequencing, or flagged with low sequencing quality. A linear mixed model was used (relmatLmer and matlm) with the kinship matrix set as a random effect. Kinship was calculated per species as the covariance matrix of the respective SNP set. A second GWAS iteration was performed for every phenotype, using the same model but adding the most significant SNP as a fixed covariate in the matlm command. This was done to help distinguish multiple peaks as well as uncover other significantly associated loci. For all GWAS the Bonferroni method was used to correct for multiple testing (0.05/Number of tested SNPs). GWAS results were processed per phenotype and species, generating Manhattan plots and summary statistics tables automatically using the script Process_GWAS_Parallel.R. Manhattan plots were generated using ggplot2 (Version 3.4.4) and cowplot (Version 1.1.1) [41, 42].

### Identification of candidate genes

QTLs were defined as the primary most significant peak of the first iteration GWAS, and any secondary significant peak of the second iteration GWAS. The candidate region was determined by the lower and upper bound SNP that was still above the Bonferroni-corrected threshold of each QTL. The functional Salinas reference genome V8 (GCF_002870075.2) annotation was used to identify genes within the candidate region, where any gene falling within a QTL interval was considered a candidate gene [43]. Functional annotation was used to prioritize candidate genes, with genes being considered primary candidates if they were supported by previous studies, homologous relationships, or functional predictions. This approach provides a practical solution to prioritize candidate genes, but it does not exclude the possibility that genes without functional annotation could be causal, nor does it eliminate the potential that the causal polymorphism lies outside of the QTL interval.

## Results

### The *Lactuca* genus exhibits broad phenotypic variation

To gain insight into the phenotypic variation in lettuce and wild *Lactuca* species we collected 76 phenotypes from the CGN *Lactuca* collection and measured 59 additional phenotypes from images available at the CGN website. This resulted in 135 total phenotypes across 550 *Lactuca* accessions (**Tables S2**+**S3**, **Figure 1A**). Species in the collection are not equally represented, with *L. sativa* (n=190) and *L. serriola* (n= 202) accessions making up the largest part included in this study, followed by *L. saligna* (n=58), *L. virosa* (n= 49) and finally the rest of the wild species (n=51) (**Figure 1B**). The collected phenotypes encompass a broad and diverse range of plant traits, represented by binary, ordinal, and quantitative data types. These traits include plant and leaf morphology and pigmentation, seed and flower characteristics, and pathogen resistance (**Figure 1A**). To explore relationships between phenotypes we perform Spearman correlation on quantitative and ordinal traits across all species (**Figure 1C**). We expect correlations between traits to be biased due to the sparse and uneven representation of species across phenotypes, however, distinct clusters of correlated traits still emerge such as morphology traits. Pathogen resistance traits form clearly correlated groups, whereas seed and flower traits are more dispersed.

**Figure 1:**
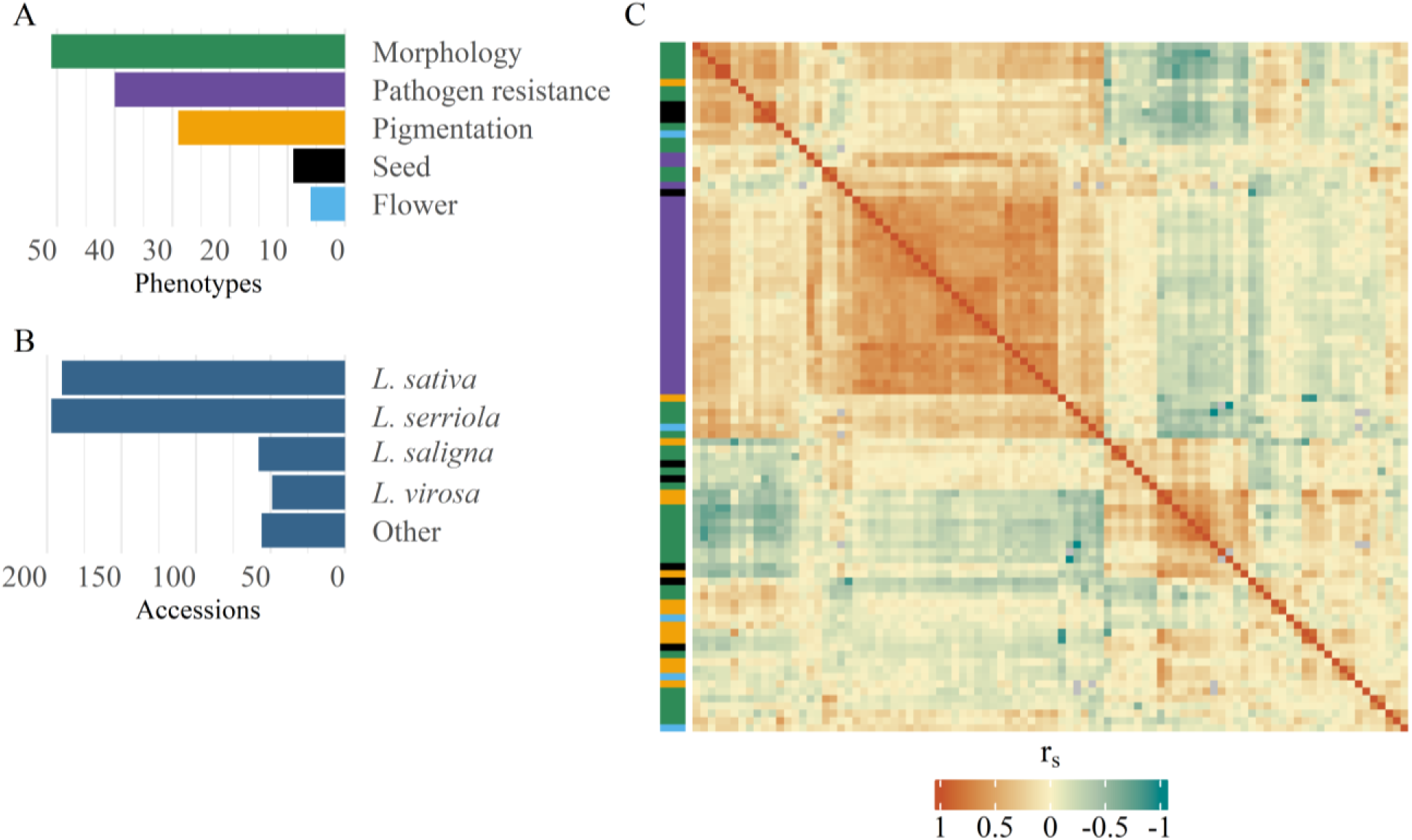
Overview of collected phenotypes across species. A) Bar graph showing the number of phenotypes per class. B) Bargraph showing the number of accessions per species. C) Heatmap of pairwise Spearman correlation of quantitative and ordinal phenotypes. Pairwise Spearman’s rho is indicated by color. Left color bar encodes phenotype classes as seen in panel A.

To assess whether traits adequately represent plant variation, we calculated weighted Gower’s distance across all, except the pathogen resistance traits, and performed hierarchical clustering. Since pathogen resistance traits are largely independent of plant morphology, we excluded them from this analysis to focus on structural and developmental traits. The resulting clusters show that accessions of the same species generally group together (**Figure 2**). Despite including a wide range of phenotypic traits, the primary split in the dendrogram appears to be driven by morphological differences (**Figure S1**). *L. saligna* accessions group with a larger subset of *L. serriola* accessions, while *L. virosa* accessions show closer clustering among themselves. *L. sativa* accessions also form a distinct cluster. This pattern is likely driven by leaf shape, as *L. virosa* and *L. sativa* exhibit more similar leaf morphology compared to *L. serriola* and *L. saligna*. This highlights that the phenotypic variation captured in our dataset effectively reflects the phenotypic diversity within *Lactuca* species included in this study.

**Figure 2:**
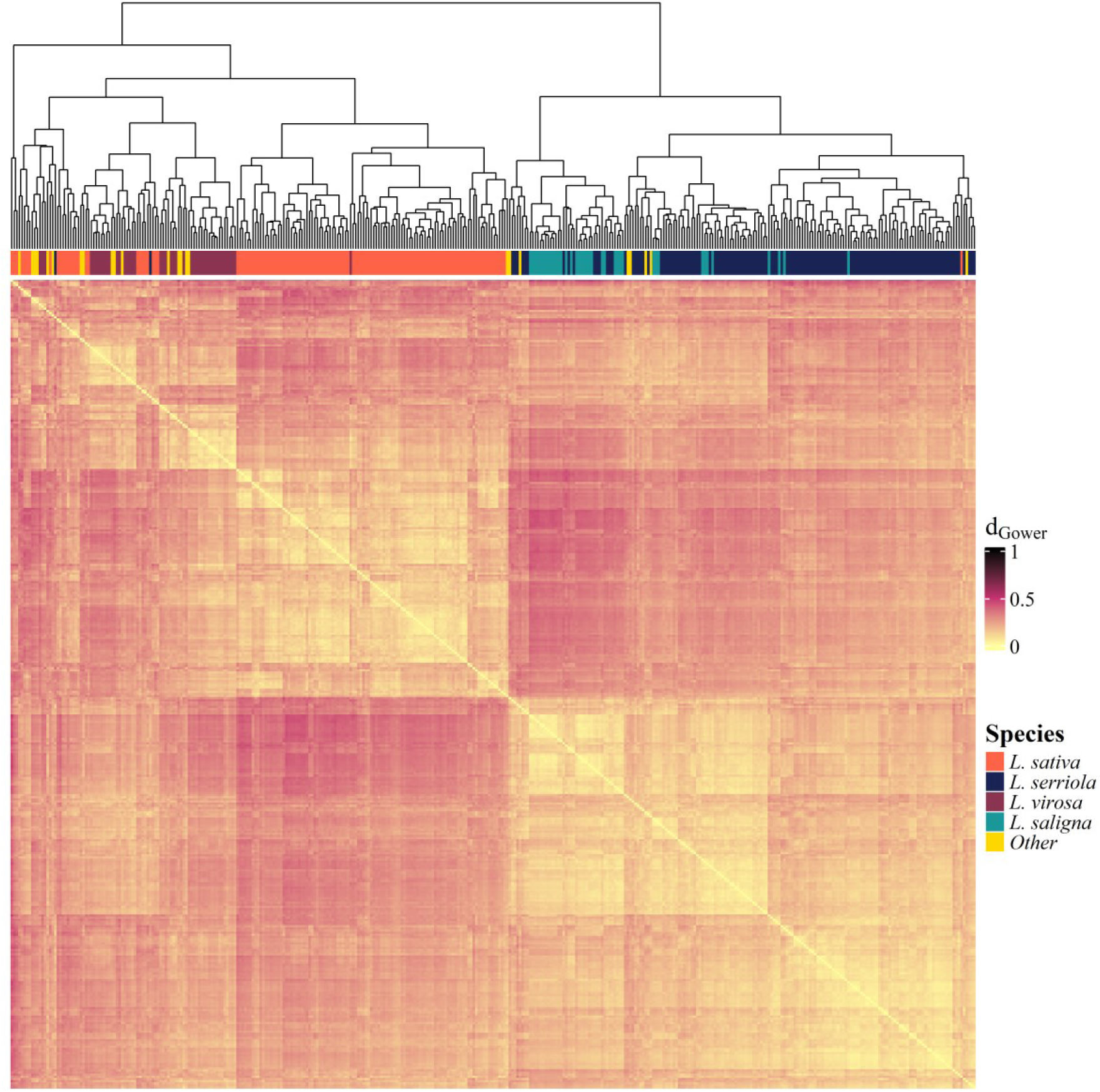
Phenotypic distance between *Lactuca* accessions. Heatmap of Gower’s dissimilarity matrix based on phenotypes between *Lactuca* accessions. d_Gower_ ranges from 0 to 1, with 0 being completely similar and 1 being completely dissimilar. The color-bar on top encodes species of respective accessions.

### Linking phenotypic variation to phylogenetic distance in *Lactuca*

Linking the phenotypic variation with the phylogenetic distance between the *Lactuca* species reveals a diverse and phenotype-specific relation with evolutionary distance, including *L. sativa* gene pools (**Figure 3**). Seed coat and flower phenotypes, as well as leaf shape are distinct phenotypes that show a broad variation across all species included in this study. Here we use the length-over-width ratio as a proxy of seed coat shape, ranging from round to elongated. The roundest seeds were found in *L. virosa, L. aculeata* and *L. georgica*, while *L. quercina* possesses very thin seed coats. Members of the primary gene pool show intermediate seed coat shapes, except for *L. aculeata*.

**Figure 3:**
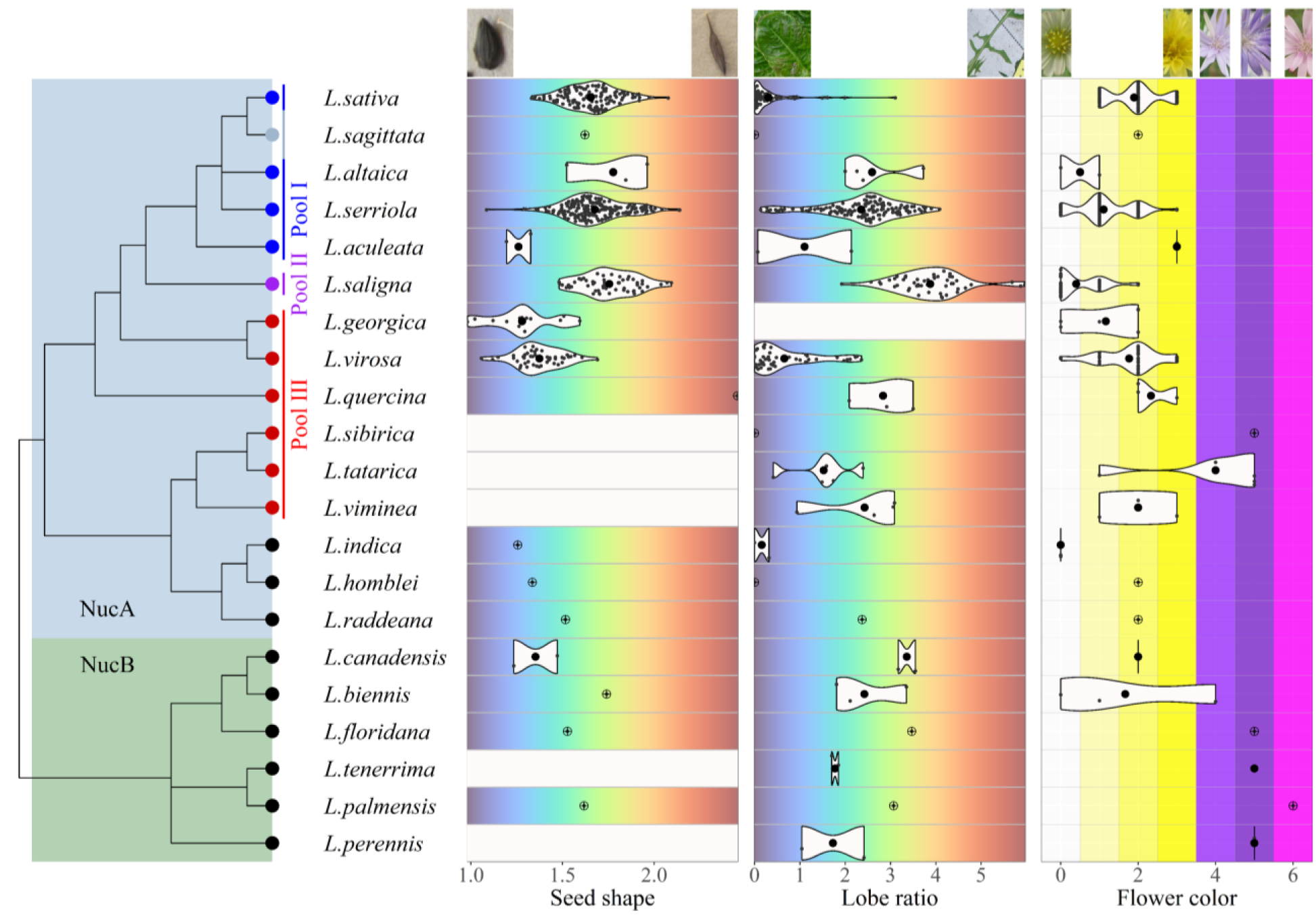
Phenotypic variation of seed, flower, and leaf traits in *Lactuca* species. Nuclear clades (from Jones *et al*. 2018) are indicated by the blue and green background square. Leaf node color indicates gene pools of L. sativa, blue: primary pool, light blue: unknown, purple: secondary pool, red: tertiary pool, black: outside the gene pool. Seed coat shape is described by the log2-transformed length-width ratio. Leaf lobing is described by the log2-transformed ratio of widest to thinnest part of a leaf lobe. Flower color is coded numerically from 0 = white to 6 = pink. Violin plots show the distribution per species, small black points indicate single observations. Crossed points indicate that only a single datapoint exists for that species. White tiles indicate missing data. Pictures at the top exemplify the phenotype based on the trait score.

Leaf lobing is a highly diverse trait across Lactuca species, ranging from weak to pronounced lobing in nearly all species. To quantitatively measure lobing we calculate the length ratio of the widest over thinnest part of a leaf lobe. *L. saligna* displays the most extreme form of leaf lobing. *L. sativa* and *L. serriola* show the greatest variation, with *L. sativa* tending towards weak or non-lobed leaves, while *L. serriola* exhibits more pronounced lobing. Additionally, *L. virosa*, *L. sibirica*, *L. indica* and *L. homblei* demonstrate a range from weak to absent lobing, contrary to other wild species.

For flower color, most species have white to yellow flowers. Light purple to pink colored flowers are only observed for species outside of the *L. sativa* gene pools, except for *L. sibirica* and *L. quercina* of the extended tertiary pool. The shades of yellow show variation between the species, with *L. serriola* and *L. saligna* having pale yellow to almost white flowers compared to *L. sativa* and *L. virosa*. Floret number across species is highly variable, ranging from 5 (*L. viminea*) and to 38 florets (*L. serriola*) (**Figure 3S**). Within the *L. sativa* gene pool, *L. viminea* and *L. saligna* had less florets on average. A loose composite flower morphology was found to be an *L. sativa* specific trait (**Figure S3**).

### GWAS in *L. sativa* and *L. serriola* reproduces known QTLs

To identify genetic loci underlying phenotypic variation in *L. sativa* and *L. serriola*, we performed two-step GWAS per species, examining 80 traits in *L. sativa* and 60 in *L. serriola* (**Table S1+S2**). Implementing a two-step GWAS approach enhanced our ability to untangle multiple QTLs per trait, identifying additional candidate loci. Across trait classes, this approach revealed both trait-specific and shared QTLs, distinguishing primary loci (first iteration) from secondary loci (second iteration). We defined shared QTLs as those co-localizing across at least two traits within the same class.

Across leaf pigmentation, pathogen resistance and leaf morphology traits, several QTLs have already been identified for *L. sativa* and *L. serriola* (**Figures S4+S5**). In this section we give an overview of the QTLs we have reproduced in each species (**Table S4)**.

Leaf pigmentation through anthocyanin accumulation in *L. sativa* is a poly-genic trait linked to several QTLs which have previously been identified. Two of those, located on chromosome 5 and 9, we have reproduced in all 11 anthocyanin-related traits included for GWAS in *L. sativa* [44–46]. Underlying the QTL on chromosome 5 at 85.5Mbp is the transcription factor *LsMYB113*, while the QTL on chromosome 9 at 152.7Mbp harbors the candidate gene *LsANS* (**Table S4**). In addition, the green leaf color intensity in *L. sativa* has previously been linked to a QTL on chromosome 4, which we reproduce as well. *LsGLK* at 101.3Mbp has been identified as the candidate gene, leading to pale green leaf color through alternative splicing [47]. Through iterative GWAS we reproduced a secondary QTL on chromosome 8 that is related to anthocyanin intensity in adult plants. This QTL has previously been identified for anthocyanin content in a GWAS population of field-grown lettuce, with the most likely candidates being a predicted MYB119 transcription factor at 105.2 Mbp, and a predicted FLAVODOXIN-LIKE QUINONE REDUCTASE 1 (*FQR1*) at 104.9Mbp [36].

Leaf morphology is a broad trait group that includes both agronomic traits specific to *L. sativa*, such as leaf margin undulation and head formation, and general leaf morphology traits like leaf division, venation patterning, and cotyledon shape. These traits have been studied extensively in *L. sativa*, with multiple QTLs identified [8, 17, 48]. GWAS on these traits confirms all already identified QTLs (**Table S4**). Additionally, for two of these traits, we analyzed image-derived phenotypes (AdLeafLobes for leaf division and AdLeafUnd for CGNLeafUndulation) using GWAS, again confirming the major QTLs. Leaf division is the only trait in this group for which a QTL has also previously been identified in *L. serriola*, which overlaps with the QTL on chromosome 3 in *L. sativa* [8]. In addition, we found that leaf tip shape and leaf form in *L. serriola* are also associated with this QTL, which has not been reported before (**Table S4**).

Pathogen resistance to *Bremia lactucae* is the most extensively researched trait in *L. sativa* and *L. serriola*. For GWAS in *L. sativa*, all 25 pathogen resistance traits exhibit multiple major QTLs, many of which overlap with previously reported loci [8, 49]. These QTLs align with known major resistance clusters (MRC) on chromosomes 1–5, 8, and 9 [49]. GWAS in *L. serriola* on *Bremia* resistance has been performed previously as well [8]. We reproduce the major QTLs for *L. serriola* on chromosomes 1,2 4 and 8 that overlap with the known MRC in *L. sativa* [49]. Additionally, we detected three novel *L. serriola* QTLs: one on chromosome 1 (47.6–50.1 Mbp), associated with resistance to *Bremia lactucae* races BL12 and BL29, as well as the aphid *Nasonovia ribisnigri* (**Table S4**), another on chromosome 2 at 8.5 Mbp (BL6, BL7, BL13, and BL27) within the MRC in *L. sativa*, and a third on chromosome 8 at 73.6 Mbp (BL12, BL18, BL20, BL22–24), with seven predicted LRR genes located 868 kb upstream (**Table S4**).

Finally, for flower-related phenotypes we reproduced QTLs for two phenotypes, petal tip anthocyanin and petal stripes. GWAS on both shows a known locus on chromosome 9 that underlies various anthocyanin traits through a point mutation in *LsANS*, including anthocyanin in the petal tips [8].

### Novel QTLs identified for pigment traits in *L. sativa* and *L. serriola*

Through iterative GWAS, we have identified novel QTLs linked to leaf pigmentation in both *L. sativa* and *L. serriola* (**Table 1**, **Figure 4**). For anthocyanin in *L. sativa*, we have identified 3 novel QTLs on chromosome 3 (132.24 Mbp), chromosome 4 (303.88 Mbp) and chromosome 9 (93.39 Mbp). The locus on chromosome 4 is approximately 40 Mbp broad, and unique to a diffuse leaf anthocyanin pattern (LAPD), harboring two strong candidate genes, a predicted MYB113 transcription factor homologous to *AtMYB113*, and a predicted chalcone synthase, homologous to *AtTT4*. Both are copies of key genes in the anthocyanin biosynthesis pathway of *L.* sativa, but have not been linked to anthocyanin variation yet, suggesting novel candidate genes controlling anthocyanin accumulation in *L. sativa* [15]. The locus on chromosome 3 at 132.24 Mbp is shared between CGN-derived traits for red leaf color (LCR) and leaf anthocyanin content (LAC), with a predicted 2-oxoglutarate (2OG) and Fe(II)-dependent oxygenase superfamily protein as the strongest candidate gene. This class of enzymes is widely represented in plants and involved in various metabolic pathways, such as the flavonoid pathway [50]. The locus on chromosome 9 is shared among the same two traits, as well as picture-derived anthocyanin intensity in adult leaves, and harbors three predicted Flavonoid 3’-monooxygenase (*F3’H*) genes as strong candidates. *F3’H* is a key enzyme in the anthocyanin biosynthetic pathway of many plants, including lettuce [44].

**Figure 4:**
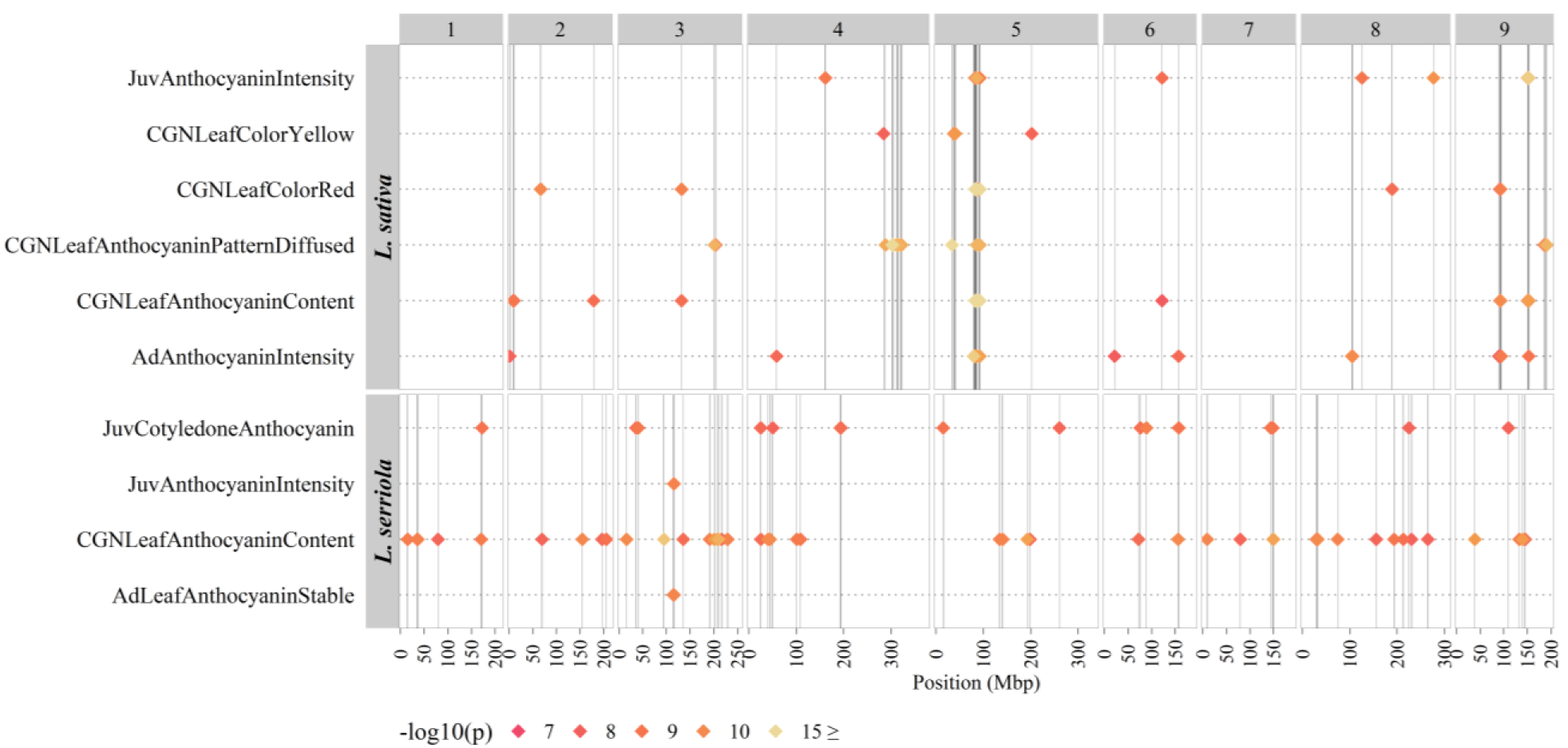
Multi-Manhattan plot of all pigment-related GWAS with novel QTLs. GWAS results summarized for all phenotypes with at least one primary significantly associated QTL in *L. sativa* and *L. serriola*. Position in mega base pairs is shown on the x-axis, chromosome numbers are indicated on top, trait and species names are shown on the y-axis. QTLs are indicated by diamonds and significance as – log10 (p) is indicated by color. Vertical lines mark the position of each QTL within a trait group.

**Table 1:**
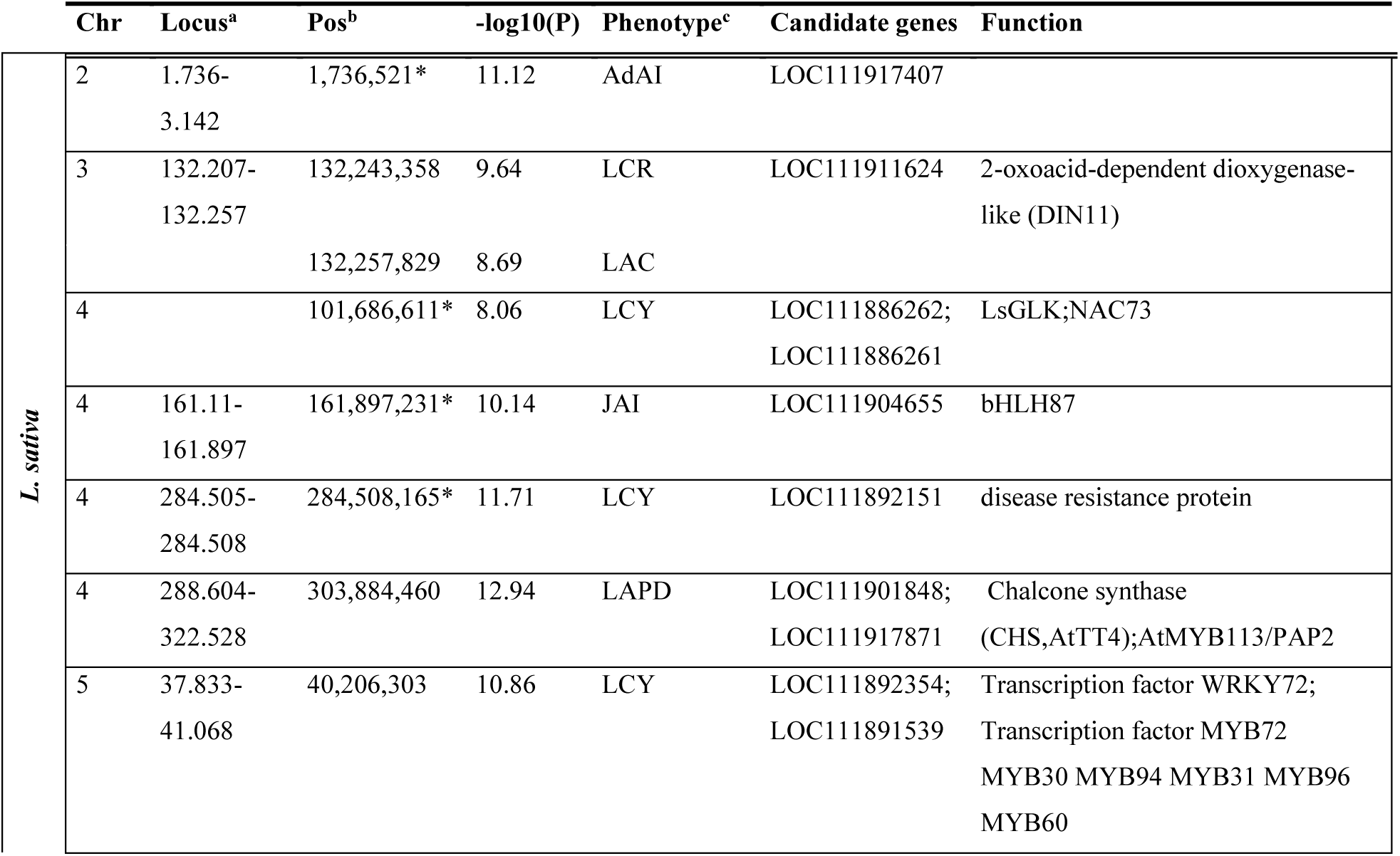
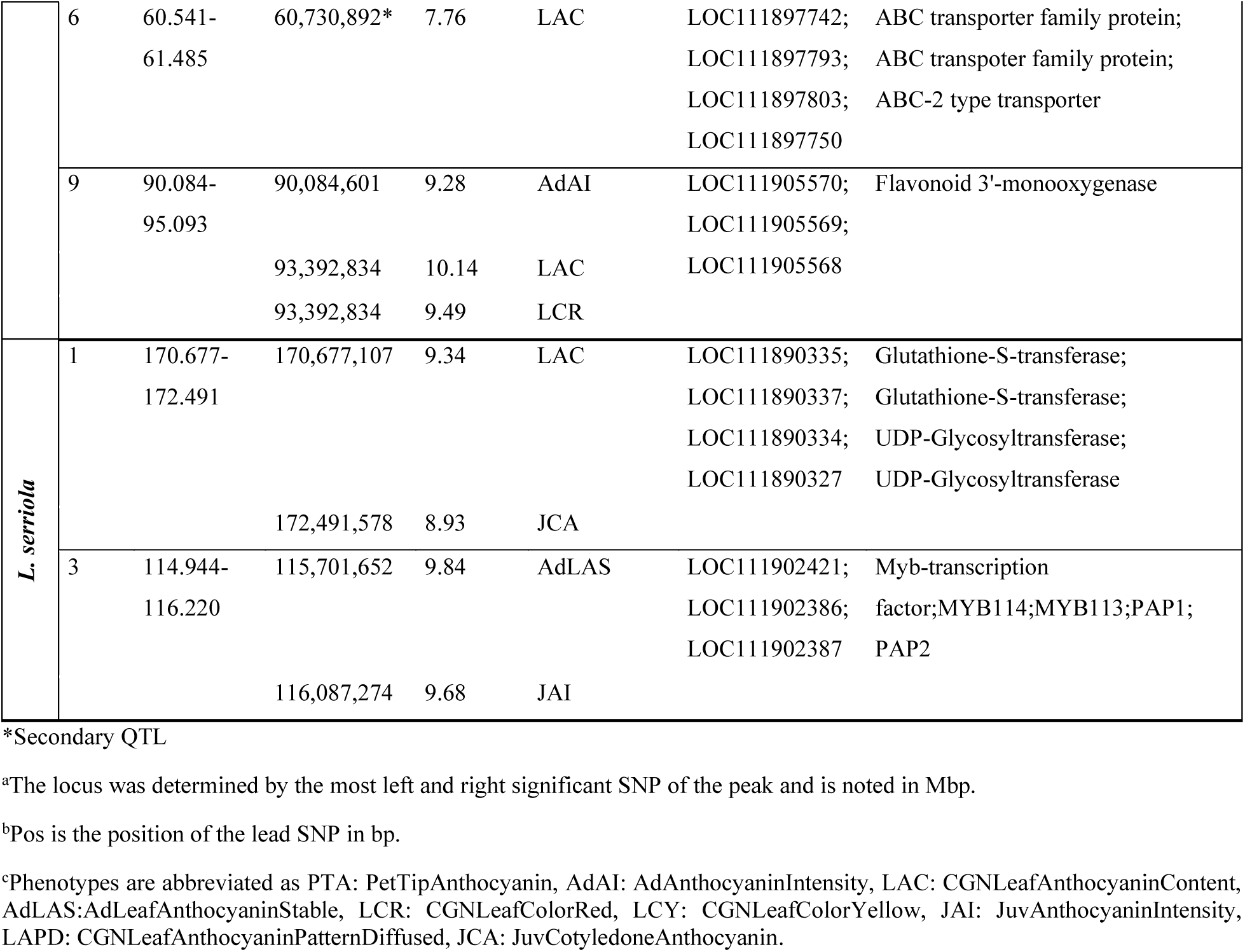
Pigment-related QTLs in *L. sativa* and *L. serriola*.

For leaf anthocyanin intensity in adult plants (AdAI) we identified a secondary QTL on chromosome 2 at 1.7 Mbp, falling within the MRC2, with no clear candidate gene present. For anthocyanin intensity in juvenile plants (JAI) we identified a secondary QTL on chromosome 4 (161.89 Mbp), with a predicted *bHLH87* transcription factor as the primary candidate gene. Anthocyanin biosynthesis in plants is controlled by a transcriptional complex, including a *bHLH* transcription factor [51]. For leaf anthocyanin content (LAC) we identified a QTL on chromosome 6 (60.73Mbp), where 3 predicted ABC transporters are located. ABC transporters are known to aid in accumulation of anthocyanin in plant tissues for various plants, making this locus a novel candidate for ABC transporters involved in anthocyanin accumulation in lettuce [43–46].

In *L. serriola*, no known QTLs exist for red leaf color traits. We identified a novel QTL on chromosome 3 at 116 Mbp, associated with anthocyanin intensity in both juvenile and adult plants (JAI, AdLAS) (**Table 1**). This locus harbors three copies of a candidate gene predicted to be homologs of *MYB113*, *MYB114*, *PAP1*, and *PAP2* in *Arabidopsis thaliana*, which are key transcription factors regulating anthocyanin biosynthesis [51]. In *L. serriola* × *L. sativa* crosses, this locus has previously been linked to antioxidant metabolism, which includes anthocyanins, with only the *L. serriola* allele of that locus contributing to metabolite variation [21]. However, this locus has not been reported for red leaf pigmentation phenotypes. We identified an additional *L. serriola*-specific QTL linked to leaf anthocyanin content (LAC) and anthocyanin presence in cotyledons (JCA) on chromosome 1 at 170.7 -172.5 Mbp. Two copies of a predicted Gluthatione-S-transferase (*GST*) and two copies of a predicted UDP-Glycosyltransferase (*UGT*) located within that QTL are strong candidate genes. *GST*s are transporters that facilitate anthocyanin accumulation into vacuoles, and *UGT*s function as modifiers of anthocyanins and precursor metabolites [52]. Neither QTL has previously been reported for *L. sativa*, highlighting their potential role in anthocyanin biosynthesis in *L. serriola* alone.

Yellowing of leaves in lettuce (LCY) can indicate environmental stress and senescence. We identify one novel QTL for *L. sativa* on chromosome 5, with predicted *MYB96* and *WRKY72* transcription factors as most likely candidate genes (**Table 1**). Both have been linked to plant stress response through abscisic acid induced leaf senescence in several plants [53–55]. Furthermore, we find clear secondary QTLs for LCY on chromosome 4 located around 101.6 and 284.5Mbp (**Table 1**). The former overlaps with a known QTL on green leaf color, where *LsGLK* has been shown to control pale leaf color [47]. The latter has not been described yet, but falls within a predicted resistance gene, suggesting a spurious association linked to the QTL underlying GLK.

### Novel QTLs identified for morphology traits in *L. sativa* and *L. serriola*

Leaf morphology is more extensively studied in *L. sativa* than in L. *serriola*. Using standard and iterative GWAS we identified novel morphology-linked QTLs for both species (**Table 2**, **Figure 5**). For elliptic leaf shape (ALE) in *L. sativa* we identified a QTL on chromosome 2 at 30.56 Mbp, with a compelling candidate gene being a predicted auxin efflux carrier protein homologous to PIN-1;3;4;7 in *A. thaliana*. PIN proteins are well described in their role of organ patterning in plants, including leaf shape [56].

**Figure 5:**
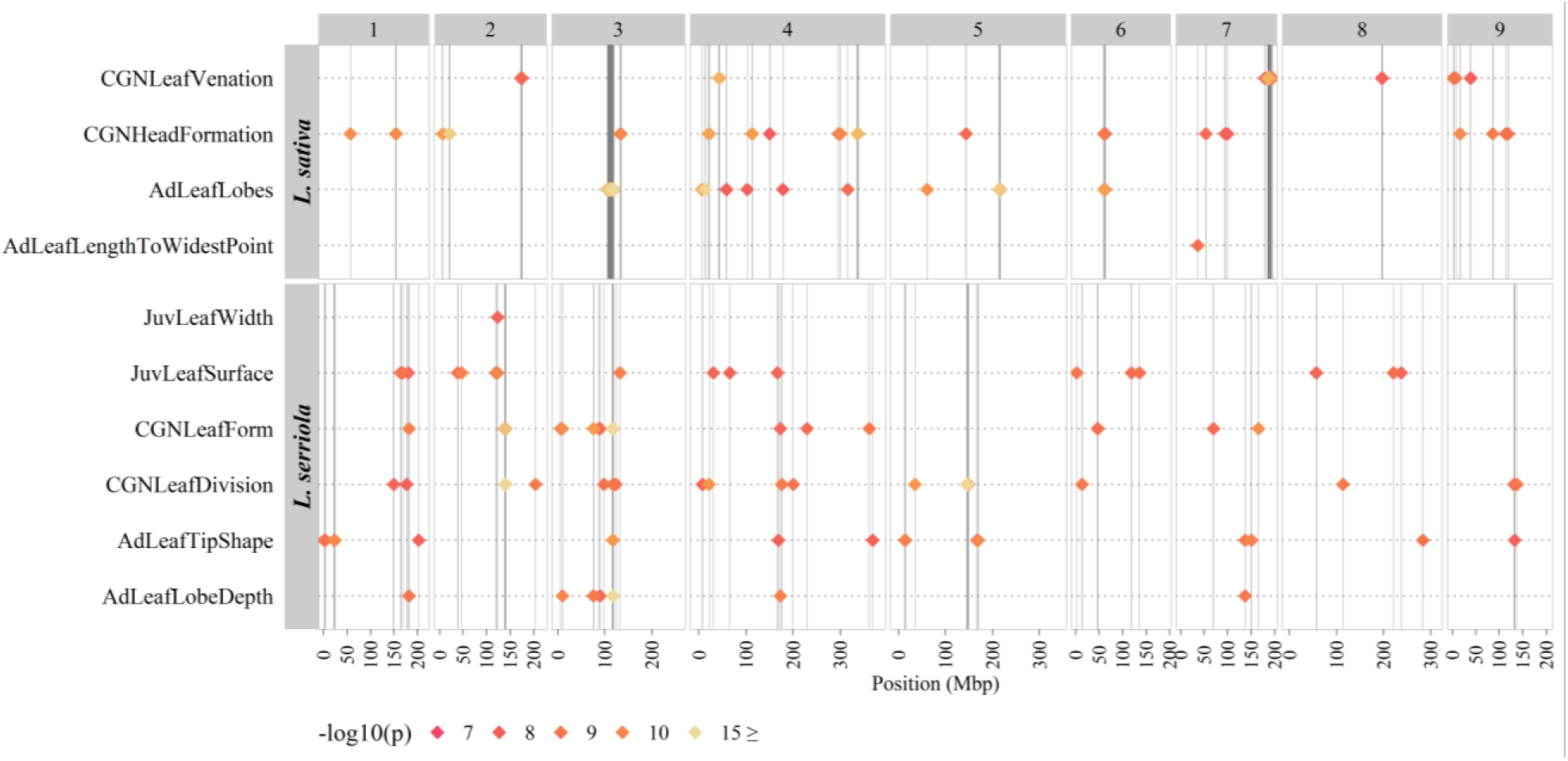
Multi-Manhattan plot of all morphology-related GWAS with novel QTLs. GWAS results summarized for all phenotypes with at least one primary significantly associated QTL in *L. sativa* and *L. serriola*. Position in mega base pairs is shown on the x-axis, chromosome numbers are indicated on top, trait and species names are shown on the y-axis. QTLs are indicated by diamonds and significance as – log10 (p) is indicated by color. Vertical lines mark the position of each QTL within a trait group.

**Table 2:**
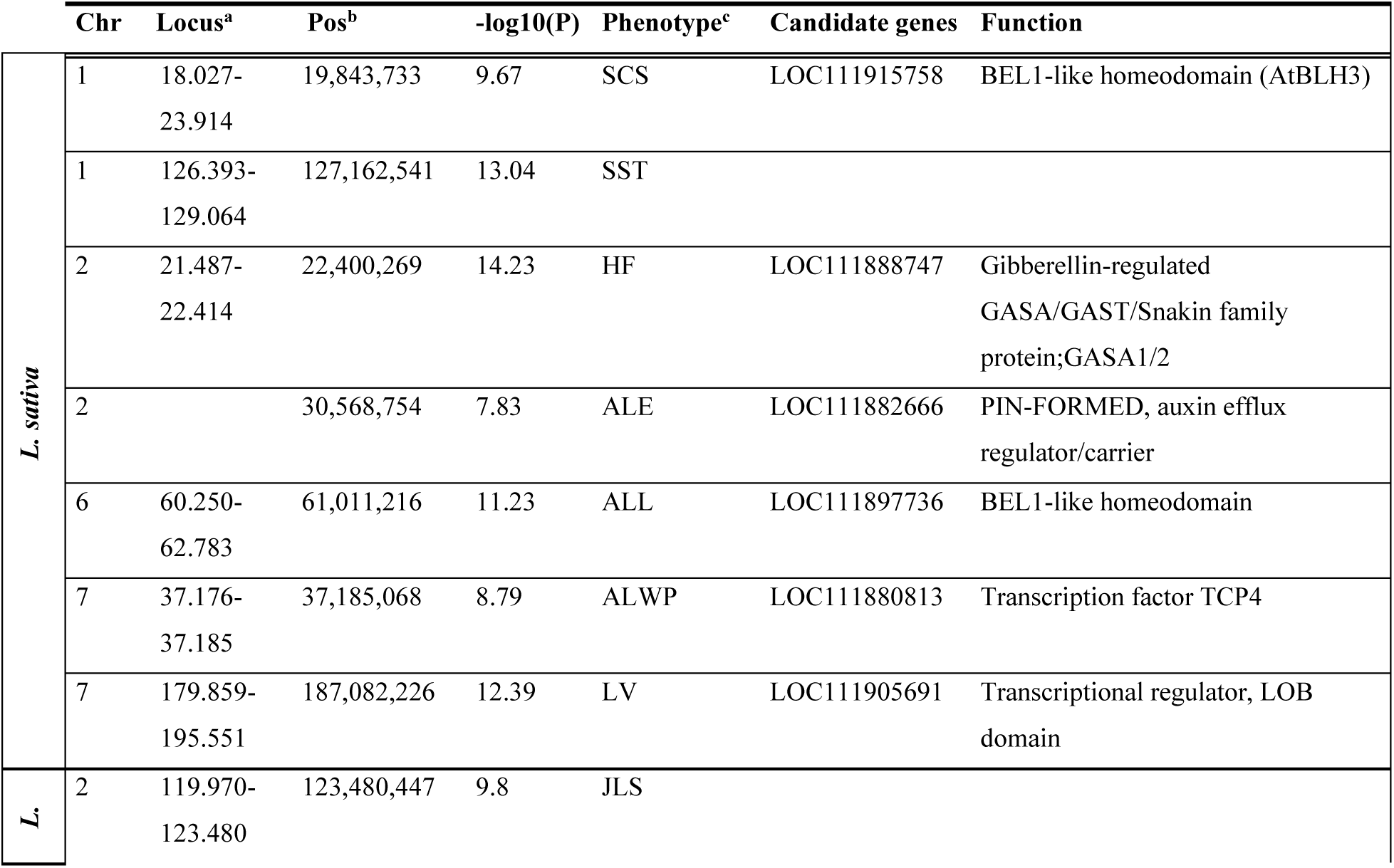
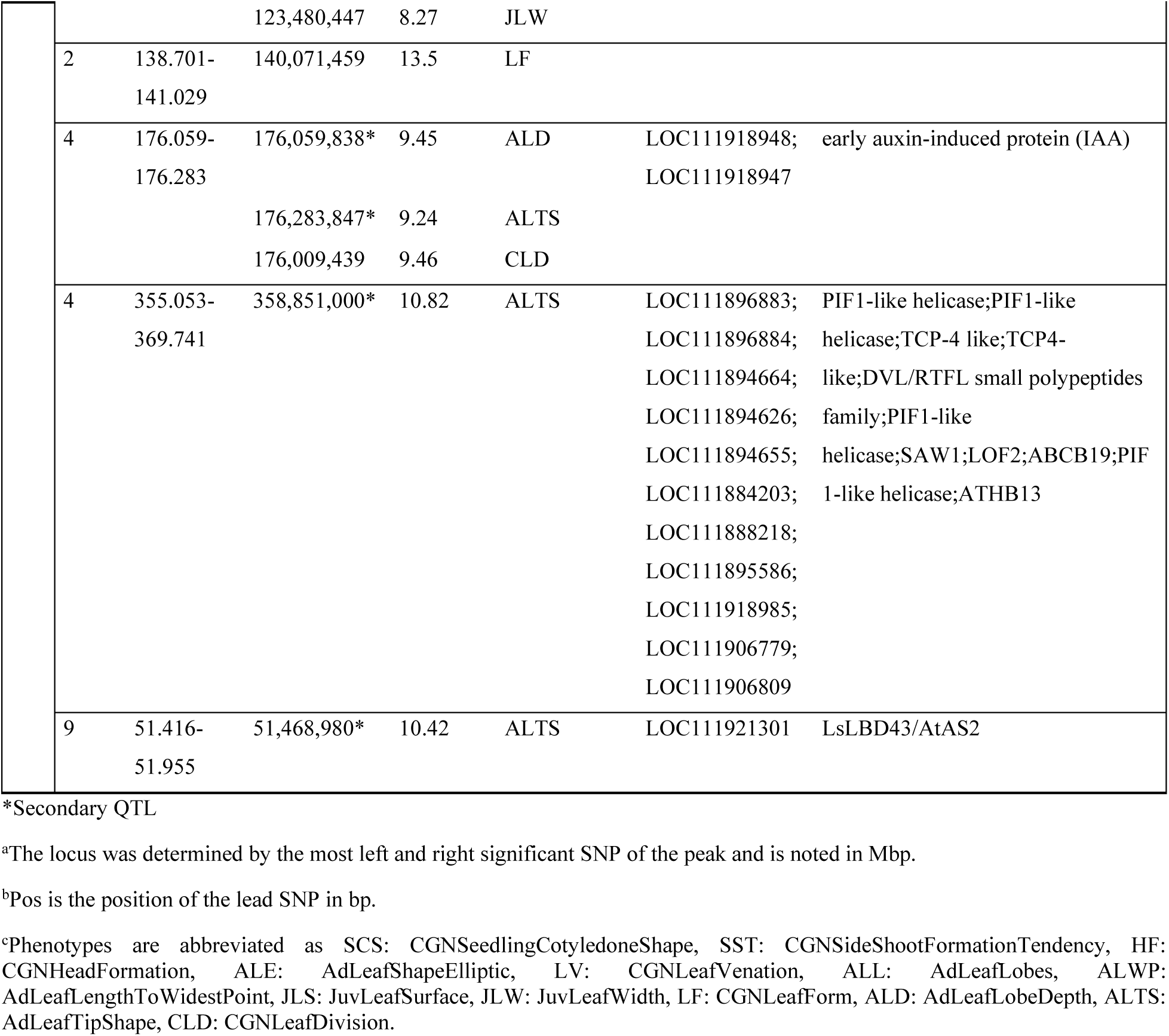
Morphology-related QTLs in *L. sativa* and *L. serriola*.

| | Chr | Locus<sup>a</sup> | Pos<sup>b</sup> | $-\log_{10}(P)$ | Phenotype<sup>c</sup> | Candidate genes | Function |
| --- | --- | --- | --- | --- | --- | --- | --- |
| *L. sativa* | 1 | 18.027-23.914 | 19,843,733 | 9.67 | SCS | LOC111915758 | BEL1-like homeodomain (AtBLH3) |
|  | 1 | 126.393-129.064 | 127,162,541 | 13.04 | SST |  |  |
|  | 2 | 21.487-22.414 | 22,400,269 | 14.23 | HF | LOC111888747 | Gibberellin-regulated GASA/GAST/Snakin family protein;GASA1/2 |
|  | 2 |  | 30,568,754 | 7.83 | ALE | LOC111882666 | PIN-FORMED, auxin efflux regulator/carrier |
|  | 6 | 60.250-62.783 | 61,011,216 | 11.23 | ALL | LOC111897736 | BEL1-like homeodomain |
|  | 7 | 37.176-37.185 | 37,185,068 | 8.79 | ALWP | LOC111880813 | Transcription factor TCP4 |
|  | 7 | 179.859-195.551 | 187,082,226 | 12.39 | LV | LOC111905691 | Transcriptional regulator, LOB domain |
| *L.* | 2 | 119.970-123.480 | 123,480,447 | 9.8 | JLS |  |  |

|  |  |  |  |  |  |  |
| --- | --- | --- | --- | --- | --- | --- |
|  |  | 123,480,447 | 8.27 | JLW |  |  |
| 2 | 138.701-141.029 | 140,071,459 | 13.5 | LF |  |  |
| 4 | 176.059-176.283 | 176,059,838* | 9.45 | ALD | LOC111918948;<br>LOC111918947 | early auxin-induced protein (IAA) |
|  |  | 176,283,847* | 9.24 | ALTS |  |  |
|  |  | 176,009,439 | 9.46 | CLD |  |  |
| 4 | 355.053-369.741 | 358,851,000* | 10.82 | ALTS | LOC111896883;<br>LOC111896884;<br>LOC111894664;<br>LOC111894626;<br>LOC111894655;<br>LOC111884203;<br>LOC111888218;<br>LOC111895586;<br>LOC111918985;<br>LOC111906779;<br>LOC111906809 | PIF1-like helicase;PIF1-like<br>helicase;TCP-4 like;TCP4-like;DVL/RTFL small polypeptides<br>family;PIF1-like<br>helicase;SAW1;LOF2;ABCB19;PIF<br>1-like helicase;ATHB13 |
| 9 | 51.416-51.955 | 51,468,980* | 10.42 | ALTS | LOC111921301 | LsLBD43/AtAS2 |

\*Secondary QTL
<sup>a</sup>The locus was determined by the most left and right significant SNP of the peak and is noted in Mbp.
<sup>b</sup>Pos is the position of the lead SNP in bp.
<sup>c</sup>Phenotypes are abbreviated as SCS: CGNSeedlingCotyledoneShape, SST: CGNSideShootFormationTendency, HF: CGNHeadFormation, ALE: AdLeafShapeElliptic, LV: CGNLeafVenation, ALL: AdLeafLobes, ALWP: AdLeafLengthToWidestPoint, JLS: JuvLeafSurface, JLW: JuvLeafWidth, LF: CGNLeafForm, ALD: AdLeafLobeDepth, ALTS: AdLeafTipShape, CLD: CGNLeafDivision.

GWAS on length-to-widest-point as description of leaf shape (ALWP) identified a novel QTL on chromosome 7 at 37.18 Mbp, with a predicted TCP transcription factor located less than 285kb from the peak. TCP transcription factors in plants have been well characterized for their involvement in cell proliferation and expansion in developing leaves [57].

Using iterative GWAS with the top SNP as a fixed effect we identified a secondary QTL for leaf lobing in adult plants (ALL) located on chromosome 6 at 61.01Mbp. This suggests an independent contribution of this locus to the phenotype. Within this locus we report on two strong candidate genes. They encode for a predicted TCP transcription factor and a BEL1-like homeobox gene (*BLH*). Several copies of *BLH*s exist in *L. sativa*, some of which have been linked to leafy head promotion; however this copy has not been characterized yet [48]. Leaf lobing in lettuce has also been linked to KNOTTED-like TALE homeobox genes (*KNOX*) [58]. In several plants *BLH*s have been shown to affect the expression of *KNOX*, causing leaf lobing and complex leaf phenotypes [59, 60]. This *BLH* gene represents a novel candidate for involvement in leaf lobing in lettuce. Interestingly, we also identify a QTL for seedling cotyledon shape (SCS) on chromosome 2 at 19.84 Mbp, with another *BLH* underlying this locus as a candidate. This suggests that at least two other distinct *BLH* copies in *L. sativa* may be involved in shaping leaf morphology at different developmental stages.

For leaf venation (LV) we identified a QTL on chromosome 7 at 187.08Mbp, with a predicted LATERAL ORGAN BOUNDARIES domain (*LBD*) gene located within this locus. Several *LBD*s have been linked to leaf morphology in *A. thaliana*, and one homolog in *L. sativa, LsLBD9* as well, however this copy (*LsLBD37*) has not been investigated further [61].

Head formation is an important agronomic trait in *L. sativa*. We report a novel QTL on chromosome 2 at 22.4 Mbp, where a predicted gibberellic acid (GA) responsive GAST1 PROTEIN HOMOLOG 1 (*AtGASA1*) is located. For heading in lettuce to occur, delayed bolting is needed, and it has been shown that the same *LsKN1* allele that is linked to head formation, is also linked to delayed bolting in lettuce as a GA biosynthesis repressor [62]. This suggests that the QTL on chromosome 2 is linked to a GA-regulated pathway that may contribute to head formation by influencing bolting timing through GA signaling.

GWAS on side-shoot formation tendency in lettuce showed a clear QTL on chromosome 1 at 127.16 Mbp. However, we could not identify a clear candidate gene.

For morphology in *L. serriola*, leaf shape traits from both CGN-derived data and image-based measurements for leaf division (CLD), leaf lobe depth (ALD), leaf form (LF) and leaf tip shape (ALTS) all share a major QTL on chromosome 3 at 117 Mbp (**Table S4**), previously linked to leaf division in *L. serriola* and *L. sativa* [7]. However, the associations of ALTS and LF with this QTL have not been reported before in *L. sativa* or *L. serriola*. Additionally, we identified a novel QTL for *L. serriola* on chromosome 4 at 176 Mbp, shared among ALD (secondary QTL), ALTS (secondary QTL), and CLD, harboring two consecutive early auxin-induced protein genes (*IAA*) as candidates, suggesting an additional locus involved in leaf morphology. These two copies have been described in *L. sativa* (*LsIAA9*, *LsIAA10*), however their relation to leaf morphology has not been investigated, also not in *L. serriola* [63]. *IAA*s have been linked to leaf patterning in several plant species through regulation of the auxin signaling pathway [64–66].

We identified additional secondary QTLs for ALTS in *L. serriola*. A QTL on chromosome 4 co-located with the known QTL for heading in *L. sativa*, where it was found that *LsSAW1* is a key regulator or maintaining leaf polarity. This broad locus is spans approx. 14Mbp, with further candidate genes present (**Table 2**). Another secondary QTL is located on chromosome 9 at 51.4Mbp. Approximately 480kb upstream of that locus we find a copy of LATERAL ORGAN BOUNDARIES DOMAIN (*LsLBD43*), homologous to ASYMMETRIC LEAVES 2 (*AtAS2*) and *LsLBD9*, suggesting another *L. serriola*-specific locus controlling leaf shape traits. In *L. sativa*, *LsLBD43* is significantly higher expressed during the adult plant stage compared to the seedling stage, which suggests a contribution of this QTL to later leaf development [61]. While iterative GWAS helps identify additional QTLs, ALTS reveals even more secondary loci. Here, we highlight the most compelling ones, but further iterations are needed to determine whether the other secondary loci represent true independent associations (**Figure S2**).

## Discussion

While phenotypic variation within *Lactuca*, particularly in *L. sativa* and *L. serriola*, is well-documented, this study provides new insights into the broader genetic and phenotypic diversity across *Lactuca* species beyond cultivated varieties. Our two-step GWAS approach identifies multiple QTLs per trait, including novel secondary QTLs, expanding the scope of those previously detected. These findings highlight the importance of understanding phenotypic variation in wild species as a first step toward identifying new breeding targets. This is particularly important when considering the potential of Crop Wild Relatives (CWR) within the *L. sativa* gene pools for breeding desirable traits like immunity and morphology. Combined with GWAS, this approach offers a more comprehensive understanding of trait genetics.

Clustering *Lactuca* accessions by morphological traits reveals species-level grouping, confirming that our approach qualitatively captures morphological differences across species. We describe a clear link between phenotypic variation and genetic differences for several traits, especially to distinguish between species. Traits like floret number, seed shape or leaf lobing highlight this, extending beyond cultivation-related traits, where only *L. sativa* stands out, like compactness of the composite flower. Combining genomic and phenotypic variation has successfully been used to distinguish duplicate *L. sativa* accessions, which are genetically identical or nearly identical samples that may have been assigned different identifiers [67]. This level of detail reinforces the validity of our approach to differentiate species, where trait differences are even more distinct.

Quantitative analysis of the phenotypic data collected in this study is challenging due to its sparsity and the presence of diverse data types, such as ordinal, continuous, binary, and categorical. Gower’s distance has been used in several studies to describe phenotypic relationships in crops, effectively integrating mixed data types and accounting for missing values [68–70]. However, it has also been shown that missing data and data type imbalances still influence Gower’s distance calculations, and consequently clustering outcomes [69, 71]. This is the inherent challenge of working with such heterogeneous datasets and requires caution when interpreting results.

Comprehensive plant image datasets, even those captured in a non-standardized manner to document germplasm collections rather than explicit phenotyping, still represent a rich resource for investigating phenotypic variation and trait genetics [72, 73]. While advances in image-based automated phenotyping are beginning to match or even exceed the accuracy of traditional manual phenotyping, achieving this level of precision requires substantial effort in training and benchmarking. Image quality, imaging methods, and the inherent complexity of plant morphology must all be carefully controlled and accounted for [74]. Until these systems are fully optimized, manual image analysis remains a valid approach, particularly when images were not taken with phenotyping as the primary goal, as is the case for images used in this study.

With GWAS in *L. sativa* and *L. serriola*, we confirmed known loci for anthocyanin biosynthesis, leaf morphology, and pathogen resistance, while also identifying novel QTLs. This is particularly noteworthy for *L. serriola*, which has been less studied through genetic mapping. For the scope of this study the two-step GWAS approach with the top SNP set as a fixed effect, provided us with a fast and automated method to detect multiple QTLs across more than 100 traits per species. Various GWAS models, such as the multi-locus mixed model (MLMM) and fixed and random model circulating probability unification (FarmCPU), employ similar but much more computationally exhaustive methodologies [75, 76].

For CGN curated traits, several have already been used for GWAS in both species, particularly by Wei *et al*. (2021). The *L. serriola* accessions in our study completely overlap with those used in their study, while our *L. sativa* dataset includes over 50 additional accessions. This expanded sample set could improve statistical power for QTL detection, but the actual gain in variation remains uncertain. Given a potential genetic bottleneck that *L. sativa* underwent during domestication, the additional accessions may not contribute significantly to overall genetic and/or phenotypic diversity [8]. We reproduced all major QTLs with a different GWAS model and SNP set, reinforcing the robustness of our and previous findings, as QTLs detected in this manner are not merely artifacts of a specific methodology, but rather likely to be genuine genetic associations. Identifying causal genetic mechanisms behind trait variation is the most labor-intensive part of GWAS or QTL mapping, requiring linkage disequilibrium analysis and additional evidence from e.g., RNA sequencing or plant transformation. Here, we report potential candidates as a starting point for further analysis, using translational evidence from other crops. Without further validation, the true causal gene(s) or variants could also be located outside the defined locus.

GWAS on the same traits between the two species makes it possible to compare the genetics underlying these traits. We find the biggest overlap in shared QTLs between the species for pathogen resistance traits, concentrated on the MRC in *L. sativa*, which has been reported before, where *L. serriola* was found to be a major donor of resistance conferring haplotypes [8, 49]. We otherwise report limited overlap of QTLs between *L. sativa* and *L. serriola* for shared traits, such as leaf lobing and leaf anthocyanin, suggesting potential differences in the genetic architecture of these traits. Phenotypic variation in plants can be redundantly controlled by paralogous genes, where the effect on the phenotype is e.g. only observed under different environmental conditions or developmental stages [77–79]. This raises questions about the extent of pathway conservation across the genus, and how the genetic bottleneck and further purifying selection in *L. sativa* may have shaped its trait genetics, purging ancestral variation, epistatic effects or redundant paralogs that control phenotype stability [6]. Leaf morphology and anthocyanin pigmentation are both complex phenotypes, with reported epistasis between several genes in *L. sativa*, like *LsMYB113* and *LsLBD9*, respectively, where we describe QTLs specific to *L. serriola* that contain paralogs of said genes.

By incorporating wild *Lactuca* species into phenotypic analyses and GWAS, we enhance our understanding of crop diversity, uncover potential breeding targets, and gain deeper insights into the genetic and evolutionary mechanisms driving phenotypic diversity across the genus. Future efforts combining multi-omics approaches and functional validation will be crucial to translate these findings into breeding strategies for more resilient and diverse lettuce cultivars.

## Supplement

**Figure S1:**
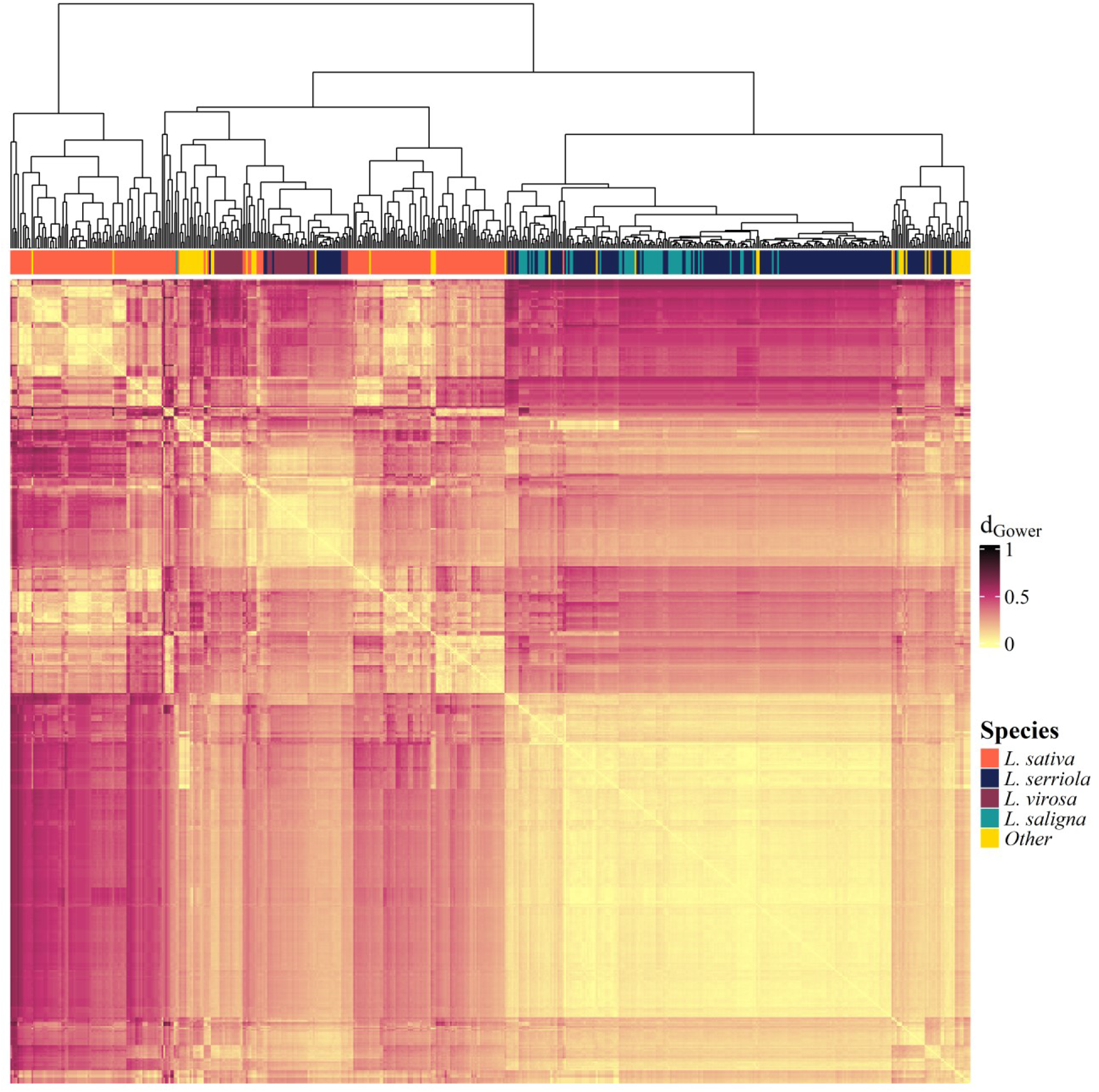
Phenotypic distance between *Lactuca* accessions. Heatmap of Gower’s dissimilarity matrix based on only morphology phenotypes between *Lactuca* accessions. d_Gower_ ranges from 0 to 1, with 0 being completely similar and 1 being completely dissimilar. The color-bar on top encodes species of respective accessions.

**Figure S2:**
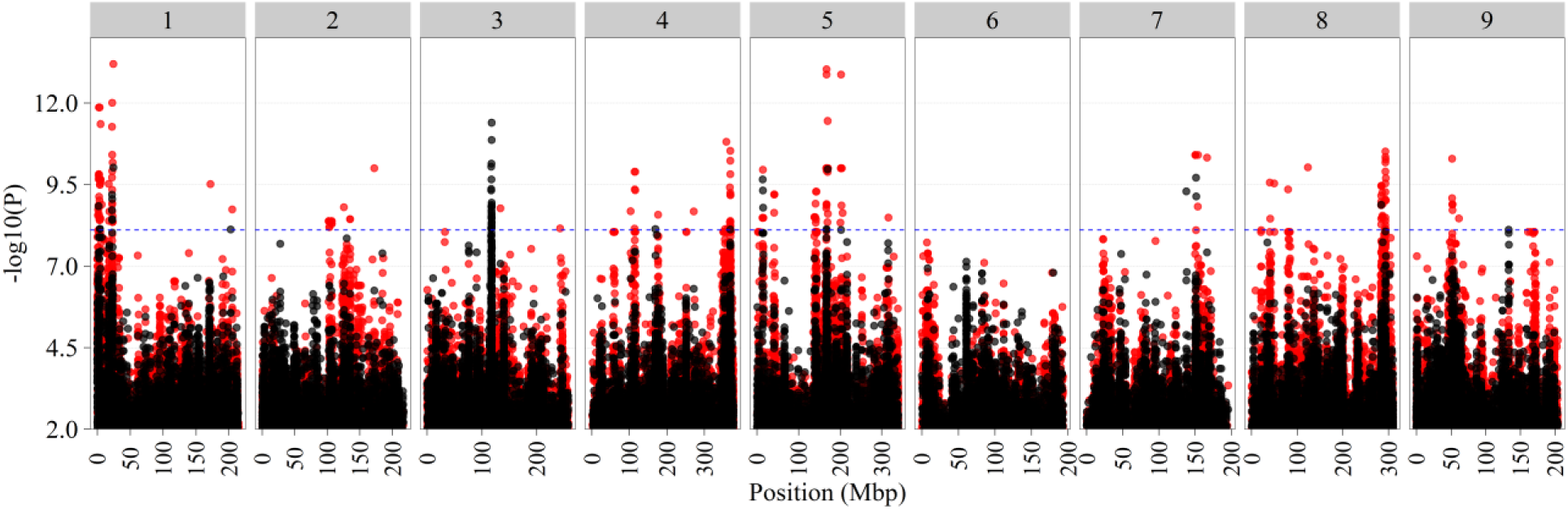
Manhattan plot of adult leaf tip shape GWAS in *L. serriola*. Two-step GWAS was performed with the top SNP on chromosome 3 set as covariate. Position in mega base pairs is shown on the x-axis, chromosome numbers are indicated on top, significance in – log10 (p) is shown on the y-axis. Bonferroni threshold of -log10(p) > 8.11 is shown by the blue horizontal line. Black dots are GWAS results without covariate, red dots are GWAS result with covariate.

**Figure S3:**
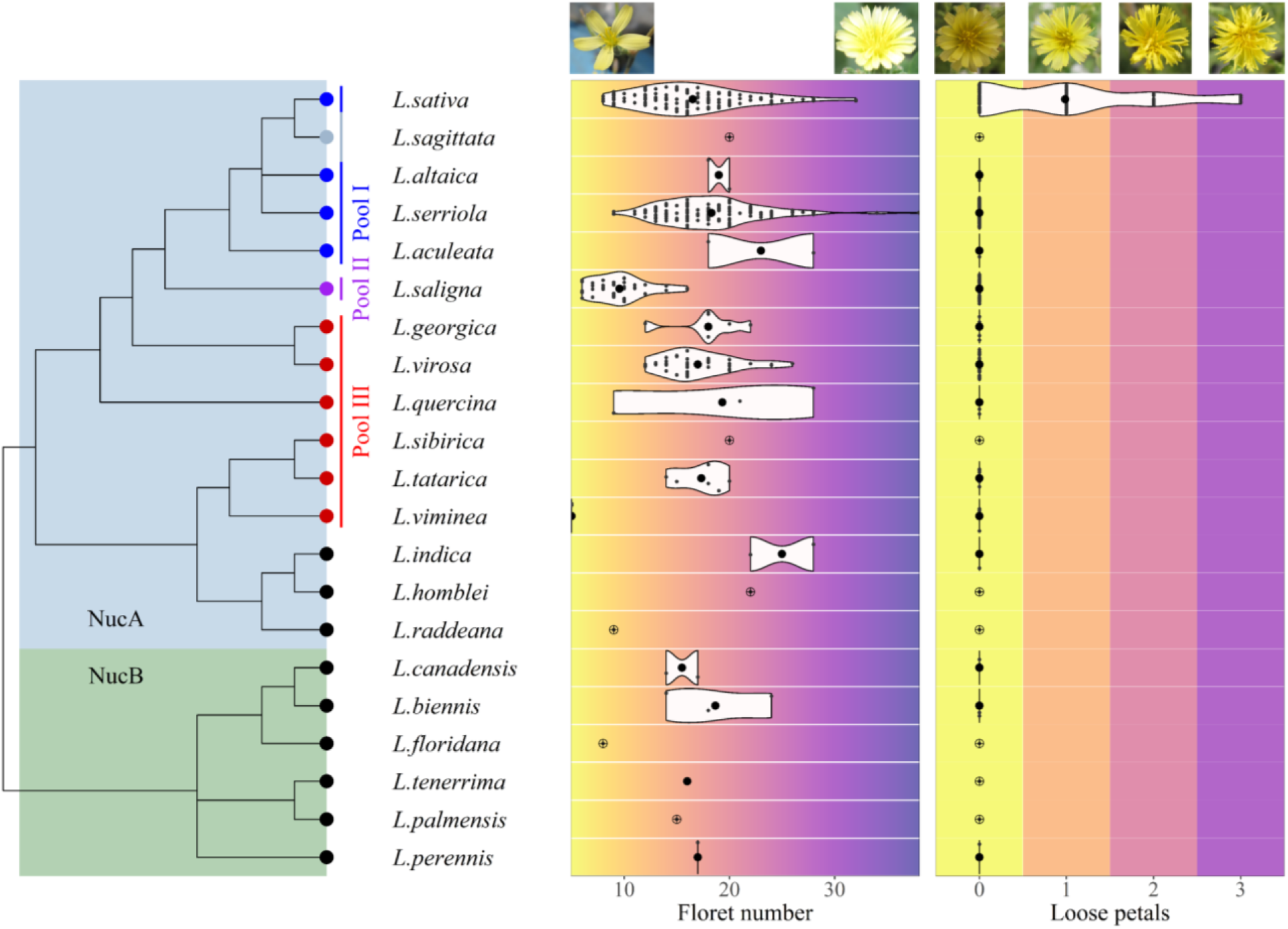
Phenotypic variation of flower traits in *Lactuca* species. Nuclear clades (from Jones et al. 2018) are indicated by the blue and green background square. Leaf node color indicates gene pools of *L. sativa*, blue: primary pool, light blue: unknown, purple: secondary pool, red: tertiary pool, black: outside the gene pool. Floret number is the count of florets in a composite flower. Loose petals describes the composite flower morphology with 0 = compact, 1 = few loose petals, 2 = intermediary loose petals, 3 = many loose petals. Violin plots show the distribution per species, small black points indicate single observations. Crossed points indicate that only a single datapoint exists for that species. Pictures at the top exemplify the phenotype based on the trait score.

**Figure S4:**
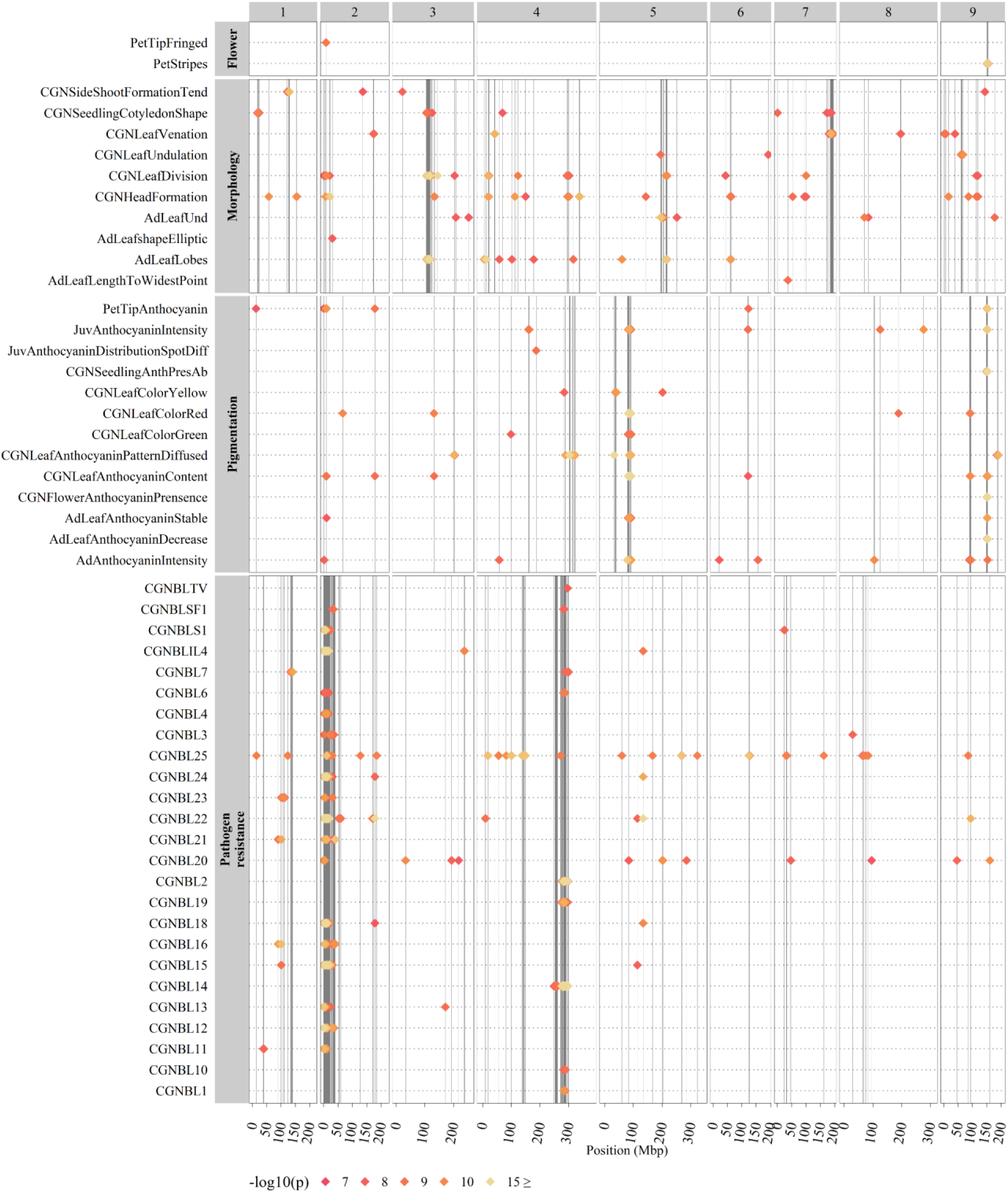
Multi-Manhattan plot of all *L. sativa* GWAS with significant QTLs. GWAS results summarized for all phenotypes with at least one primary significantly associated QTL in *L. sativa*. Position in mega base pairs is shown on the x-axis, chromosome numbers are indicated on top, trait names and trait groups are shown on the y-axis. QTLs are indicated by diamonds and significance as – log10 (p) is indicated by color. Vertical lines mark the position of each QTL within a trait group.

**Figure S5:**
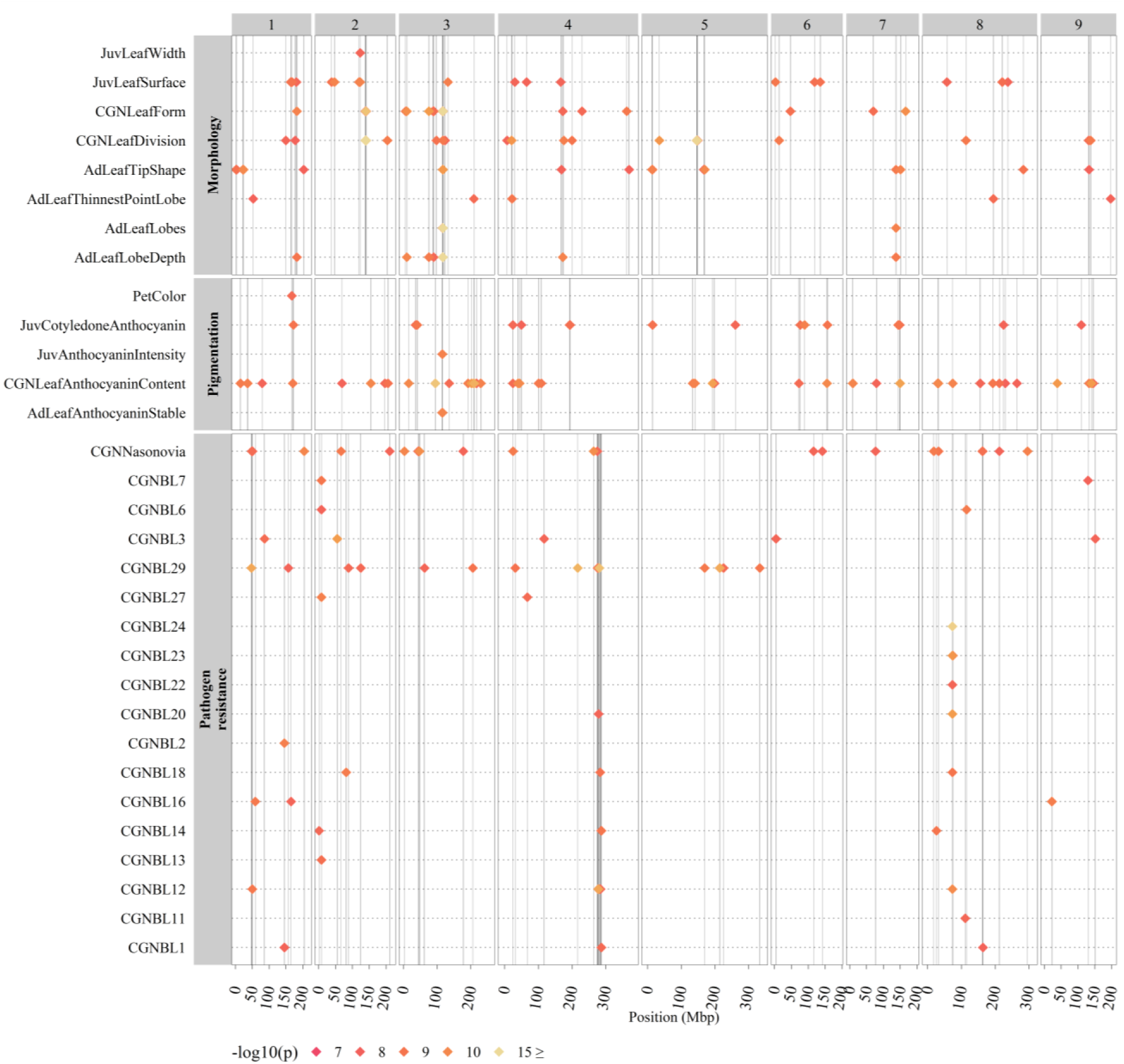
Multi-Manhattan plot of all *L. serriola* GWAS with significant QTLs. GWAS results summarized for all phenotypes with at least one primary significantly associated QTL in *L. serriola*. Position in mega base pairs is shown on the x-axis, chromosome numbers are indicated on top, trait names and trait groups are shown on the y-axis. QTLs are indicated by diamonds and significance as – log10 (p) is indicated by color. Vertical lines mark the position of each QTL within a trait group.

## Code and data availability

Scripts used for this study are available on GitHub at https://github.com/SnoekLab/ Mehrem_etal_2025_PhenoLac.

SNP data for *L. serriola* accessions is available upon request.

## Acknowledgements

We would like to thank Tom Schermer for his support during manual phenotyping. We thank the CGN for sharing plant images and phenotypic scores.

## Funding

This publication is part of the LettuceKnow project (with project number 1.2 of the research Perspective Program P19-17 which is (partly) financed by the Dutch Research Council (NWO) and the breeding companies BASF, Bejo Zaden B.V., Limagrain, Enza Zaden Research & Development B.V., Rijk Zwaan Breeding B.V., Syngenta Seeds B.V., and Takii and Company Ltd.

## Author contributions

SM, BLS and GvdA devised the study. SM and BLS analyzed the data and wrote the manuscript with input from all co-authors.

## Notes

https://github.com/SnoekLab/Mehrem_etal_2025_PhenoLac

### Competing Interest Statement

The authors have declared no competing interest.

